# Conservation of glutathione S-transferase mRNA and protein sequences similar to human and horse Alpha class GST A3-3 across dog, goat, and opossum species

**DOI:** 10.1101/2020.11.25.396168

**Authors:** Shawna M. Hubert, Paul B. Samollow, Helena Lindström, Bengt Mannervik, Nancy H. Ing

## Abstract

Recently, the glutathione S-transferase A3-3 (GST A3-3) homodimeric enzyme was identified as the most efficient enzyme that catalyzes isomerization of the precursors of testosterone, estradiol, and progesterone in the gonads of humans and horses. However, the presence of GST A3-3 orthologs with equally high ketosteroid isomerase activity has not been verified in other mammalian species, even though pig and cattle homologs have been cloned and studied. Identifying *GSTA3* genes is a challenge because of multiple *GSTA* gene duplications (12 in the human genome), so few genomes have a corresponding *GSTA3* gene annotated. To improve our understanding of *GSTA3* gene products and their functions across diverse mammalian species, we cloned homologs of the horse and human *GSTA3* mRNAs from the testes of a dog, goat, and gray short-tailed opossum, with those current genomes lacking *GSTA3* gene annotations. The resultant novel *GSTA3* mRNA and inferred protein sequences had a high level of conservation with human *GSTA3* mRNA and protein sequences (≥ 70% and ≥ 64% identities, respectively). Sequence conservation was also apparent for the 13 residues of the “H-site” in the 222 amino acid GSTA3 protein that is known to interact with the steroid substrates. Modeling predicted that the dog GSTA3-3 is a more active ketosteroid isomerase than the goat or opossum enzymes. Our results help us understand the active sites of mammalian GST A3-3 enzymes, and their inhibitors may be useful for reducing steroidogenesis for medical purposes, such as fertility control or treatment of steroid-dependent diseases.

## 1. Introduction

It is widely appreciated that reproduction of animal species important to man centers around sex steroid production by the gonads. While driven by peptide hormones from the hypothalamic/pituitary axis, production of testosterone by the Leydig cells of the testis and estradiol and progesterone by the granulosa cells and corpora lutea of the ovary are required for successful production of offspring [1–3]. Testosterone, estradiol and progesterone act in reproductive tissues by binding specific receptors which regulate the expression of sets of genes [4,5].

Testosterone, estradiol and progesterone are produced from cholesterol by a series of enzymatic modifications [6,7]. This reaction sequence differs among species: rodents utilize the Δ^4^ pathway whereas humans and other larger mammals utilize the Δ^5^ pathway [7]. The biosynthetic pathway for testosterone in human and horse testes involves the alpha class glutathione S-transferase A3-3 (GST A3-3) as a ketosteroid isomerase that acts as a homodimer to convert Δ^5^-androstene-3,17-dione to Δ^4^-androstene-3,17-dione, the immediate precursor of testosterone [8–15]. The recombinant human GST A3-3 enzyme is 230 times more efficient at this isomerization than hydroxy-Δ^5^-steroid dehydrogenase, 3β-hydroxysteroid delta-isomerase 2 (HSD3B2), the enzyme to which this activity previously had been attributed [2]. Estradiol synthesis is likewise dependent upon GSTA3-3 in humans and horses because testosterone is further converted to estradiol by aromatase [7]. GST A3-3 also participates in the synthesis of progesterone in female mammals utilizing the Δ^5^ pathway by isomerizing Δ^5^-pregnene-3,20-dione to Δ^4^-pregnene-3,20-dione (i.e., progesterone).

From a general perspective, the GST family of enzymes is recognized for having active roles in detoxification. The GSTA class of genes is best characterized in humans. It is represented by 12 genes and pseudogenes with almost indistinguishable sequences. These arose from gene duplications and in humans are clustered in a 282 kbp region on chromosome 6 [16]. Because of the redundancy of GSTA genes, the *GSTA3* genes are inexactly identified, if identified at all, in the current annotations of the genomes of most mammals. This gap in knowledge limits studies of *GSTA3* gene expression because standard *GSTA3* mRNA and protein detection reagents crossreact with other four gene products in the GSTA family, which are ubiquitously expressed [11].

Understanding the regulation of expression of *GSTA3* gene and the activities of GSTA3-3 enzyme is critical to fully comprehending the synthesis of sex steroid hormones that direct reproductive biology in animals. In extracts of a steroidogenic cell line, two specific inhibitors of GSTA3-3 (ethacrynic acid and tributyltin acetate) decreased isomerization of delta-5 androstenedione with very low half maximal inhibitory concentrations (IC_50_) values of 2.5 uM and 0.09 uM, respectively [17]. In addition, transfection of such a cell line with two different small interfering RNAs both decreased progesterone synthesis by 26%. In stallions treated with dexamethasone, a glucocorticoid drug used to treat inflammation, testicular levels of *GSTA3* mRNAs decreased by 50% at 12 hours post-injection, concurrent with a 94% reduction in serum testosterone [18,19]. In stallion testes and cultured Leydig cells, dexamethasone also decreased concentrations of steroidogenic factor 1 (SF1) mRNA [20]. In a human genome-wide study of the genes regulated by SF1 protein, chromatin immunoprecipitation determined that the SF1 protein bound to the *GSTA3* promoter to up-regulate the gene [13]. Notably, the SF1 transcription factor up-regulates the expression of several gene products involved in steroidogenesis [2]. These data indicate that there are reagents and pathways by which *GSTA3* gene expression and steroidogenesis can be altered *in vivo*. This could be useful for reducing steroidogenesis for fertility control or other medical purposes.

The equine GST A3-3 enzyme is structurally similar to the human enzyme and matches or exceeds the very high steroid isomerase activity of the latter [15]. Remarkably, the cloning of GSTA3 homologs from the gonads of domestic pigs (*Sus scrofa*) and cattle (*Bos taurus*) yielded enzymes that were much less active ketosteroid isomerases for reasons yet to be determined. In the current work, *GSTA3* mRNA was cloned from the testes of the dog and goat (eutherian mammals), and gray short-tailed opossum (metatherian mammal). The purposes were (1) to compare mRNA and protein sequences with the human and horse counterparts to search for more highly active ketosteroid isomerases, (2) to generate GSTA3-specific reagents for future studies of the regulation of *GSTA3* genes, and (3) to understand the scope of the GSTA3-3 function in steroidogenesis across a range of mammalian species in which reproduction is important to humans.

## 2. Experimental

### 2.1. Testis tissue samples and RNA preparation

All animal procedures were approved by the Institutional Animal Care and Use Committee. Adult testes used in this study were obtained from Texas A&M University owned research animals. Specifically, testes were from a large breed hound (*Canis lupus familiaris*), a goat (*Capra hircus),* and, a marsupial mammal, a gray short-tailed opossum (*Monodelphis domestica*).

Total cellular RNA was extracted from each testis with the Tripure reagent (Sigma-Aldrich; St. Louis, MO) protocol. The concentration and purity of the RNA was assessed using a Nanodrop spectrophotometer.

### 2.2. Reverse transcription

Reverse transcription reactions (25 μl) for the production of complimentary DNA (cDNA) were run with each RNA sample (500 ng) using random octamer primers (312 ng), dT_20_ primers (625 ng), dNTPs, DTT, first strand buffer, Superasin and Superscript II Reverse Transcriptase (RT), following the manufacturer’s instructions (ThermoFisher Scientific; Waltham, MA). The RT reactions incubated in an air incubator at 42 °C for three hours. The RT enzyme was inactivated by incubating at 70 °C for 15 minutes.

### 2.3. Cloning GSTA3 cDNAs into the vector pCR2.1

Nested polymerase chain reaction (PCR) was utilized to selectively amplify the *GSTA3* mRNA targets because it increases the sensitivity and fidelity of the amplification (Fig. 1). Primers were designed from BLAST analyses with human and horse *GSTA3* mRNA sequences (GenBank accession numbers **<u>NM_000847.1</u>** and **<u>KC512384.1</u>**, respectively; Table 1) to identify related dog, goat and opossum sequences for cloning cDNAs into the general use vector pCR2.1 (TA Cloning Kit; ThermoFisher Scientific). The primers were synthesized by Integrated DNA Technologies (Skokie, IL). PCR reactions (50 μl) were performed with Ex Taq enzyme (Takara Bio Inc.; Mountain View, CA) in a GeneAmp PCR System 9600 (PerkinElmer; Norwalk, CT). Primary PCR reactions (50 μl) were set up with outside primers and 1 μl of the appropriate RT reaction. Cycling parameters were 30 cycles of 94 °C for 15 seconds, 45 °C annealing for 30 seconds, and 72 °C for one minute, followed by a one-time hold at 72 °C for five minutes. Secondary PCR reactions (50 μl) were set up with inside primers and 5 μl of the primary PCR reactions as a cDNA template source instead of using the RT reaction. Cycling parameters were maintained for the secondary PCR. The PCR products were run on both a 1% agarose gel and a 0.8% low melting temperature agarose gel. Ethidium bromide staining of DNA bands was visualized under UV light. This procedure confirmed the presence of the intended PCR products of 670 to 700 base pairs in all of the secondary PCR reactions.

**Table 1.**
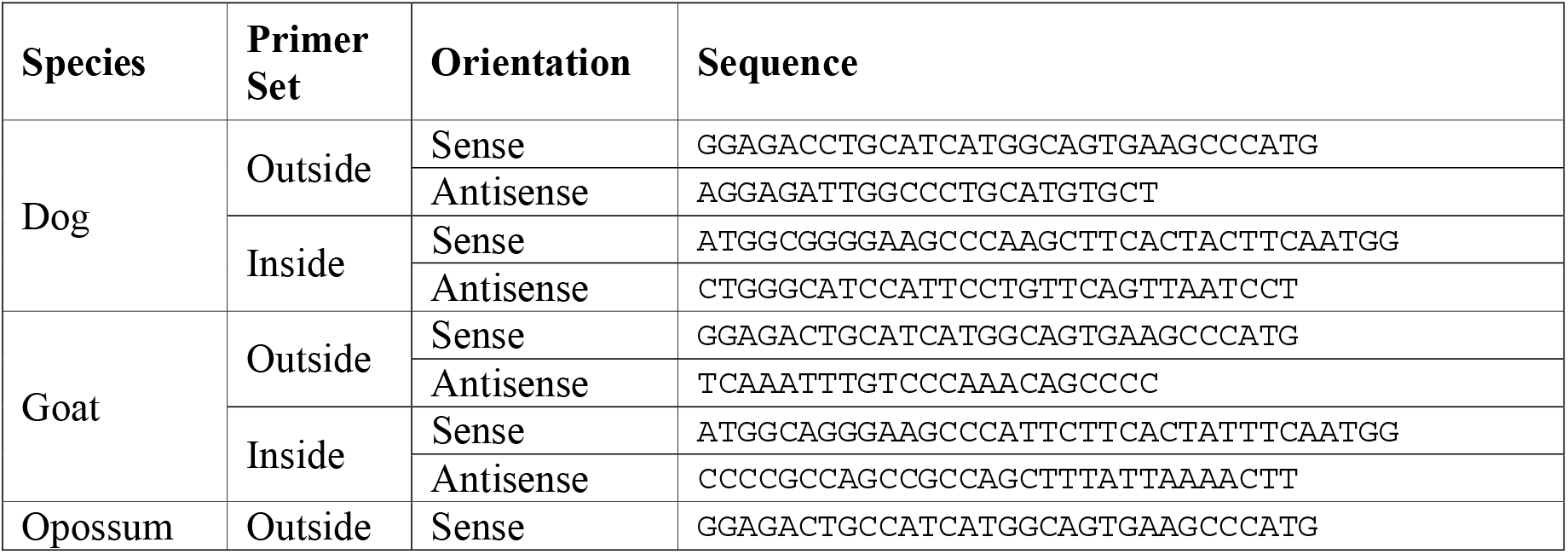
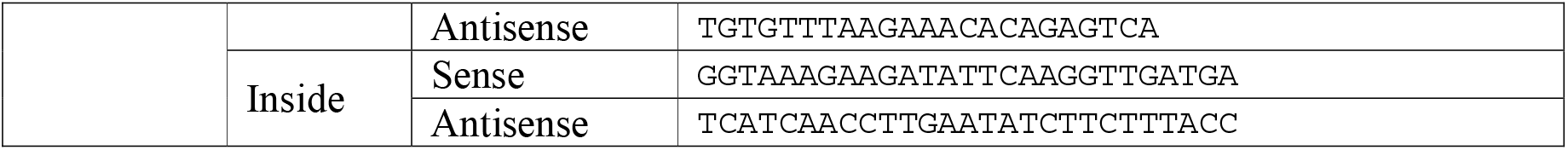
PCR primer sequences (5’-3’) for TA cloning GSTA3 cDNAs into pCR2.1.

**Figure 1.**
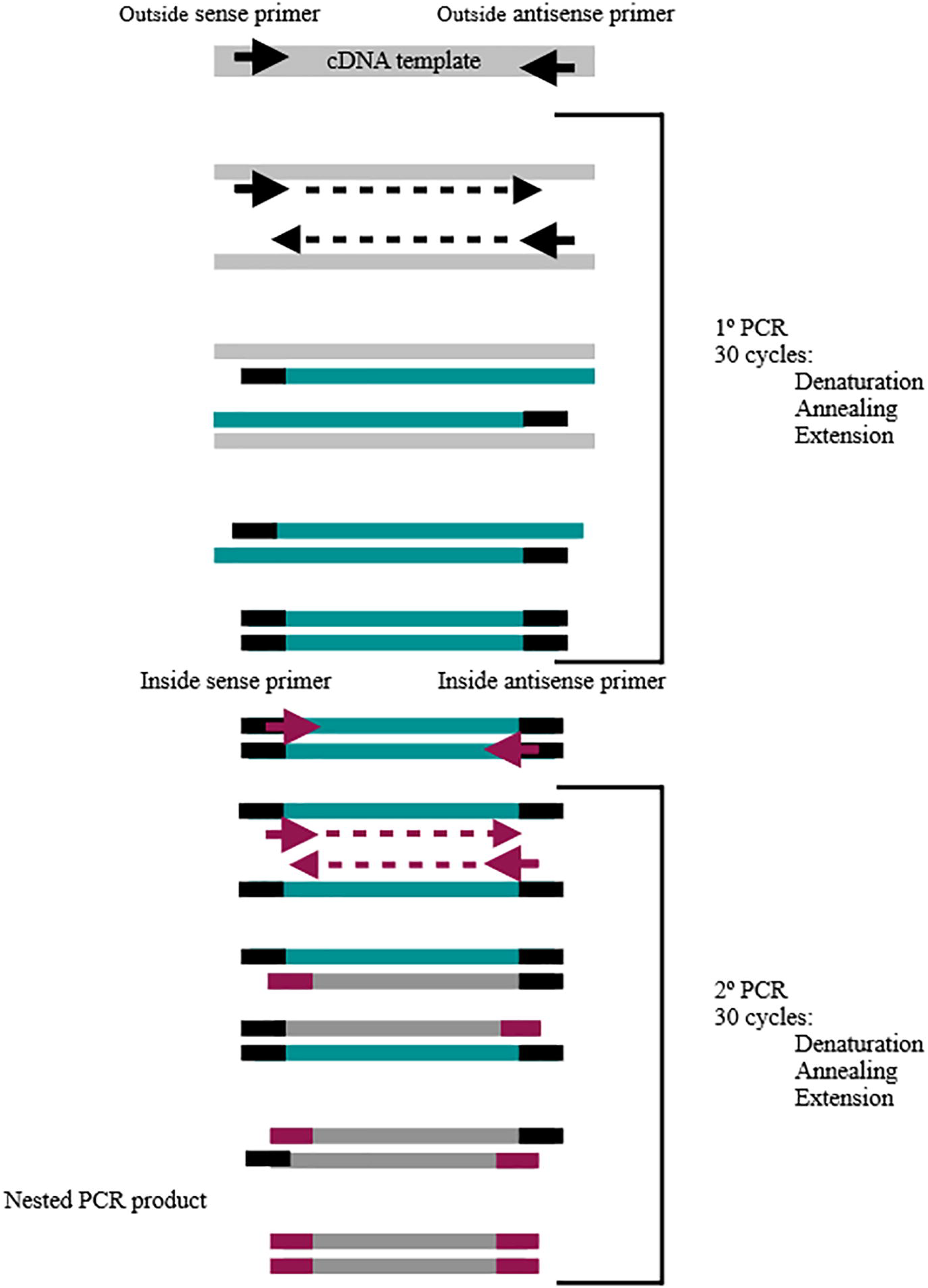
Nested PCR for amplification of GSTA3 mRNA targets.

*GSTA3* cDNA bands were cut from the 0.8% low melting temperature agarose and melted at 70 °C for 10 minutes. Ligation of the cDNA inserts to pCR2.1 was performed according to the kit’s instructions (TA Cloning kit; ThermoFisher). These ligation reactions were then utilized in transformation of *E. coli* competent cells. After incubation in SOC broth for 1 h at 37 °C, transformations were plated on LB-ampicillin agar containing isopropyl β-D-1-thiogalactopyranoside and 5-bromo-4-chloro-3-indolyl-β-D-galactopyranoside for blue-white colony selection. Plates were incubated overnight at 37 °C. Selected white colonies were cultured overnight in LB-ampicillin broth and plasmid DNA was purified using the Qiagen QIAprep Spin Kit (Germantown, MD). *GSTA3* cDNAs were released from the pCR2.1 plasmids by digestion with EcoRI before being analyzed by gel electrophoresis. Plasmid samples that demonstrated the expected 670 to 700 base pair bands were sequenced.

### 2.4. Sequencing and analysis

The *GSTA3* cDNAs in the pCR2.1 clones were amplified with PCR through the use of Big-Dye mix (Applied Biosystems, ThermoFisher Scientific) using the M13 forward and M13 reverse primers (5’-TGTAAAACGACGGCCAGT-3’ and 5’-CAGGAAACAGCTATGAC-3’, respectively). After PCR amplification, G-50 spin columns were used to remove unincorporated nucleotides. Samples were sequenced at the Texas A&M University Gene Technologies Laboratory on a PRISM 3100 Genetic Analyzer mix (Applied Biosystems, ThermoFisher Scientific). Each sequence was then compared to the NCBI GenBank database records for the human and horse *GSTA3* mRNA sequences (GenBank accession numbers **<u>NM_000847.1</u>** and **<u>KC512384.1</u>**, respectively). The basic local alignment search tool (BLAST) on the NCBI website was also utilized to check for alignments to reference mRNA sequences of other species. For each species, the sequence with the highest identity to the NCBI *GSTA3* mRNA reference sequences was translated to its amino acid sequence through the use of the ExPASy translate tool (https://web.expasy.org/translate/). Sequence validation of *GSTA3* mRNAs and proteins was obtained by locating the five amino acid residues that have been identified as key to the activity of the human GST A3-3 enzyme [9,21]. Following this analysis, *GSTA3* mRNA sequences were submitted to NCBI GenBank. Accession numbers for these sequences are given in the Results and Discussion section.

### 2.5. Cloning GSTA3 cDNA in the bacterial expression vector pET-21a(+)

Another set of optimized primers was created for cloning into the bacterial expression vector pET-21a(+) (EMD Millipore; Billerica, MA). For directional in-frame cloning into pET-21a(+), the *Eco* RI (GAATTC) and *Xho*I (CTCGAG) restriction sites were added to the 5’ ends of the inside sense and antisense primers, respectively (Table 2). Nested PCR was performed as described in section 2.3. Secondary PCR samples along with the pET-21a(+) vector were restricted with *Eco* RI and *Xho*I endonucleases. Purified bands from electrophoresis on an 0.8% LMT agarose gel were ligated. After transformation, colony growth in broth, and plasmid mini-preparation, the plasmid cDNAs were sequenced with the T7 promoter and the T7 terminator primers (5’-TAATACGACTCACTATAG-3’ and 5’-GCTAGTTATTGCTCAGCGG-3’, respectively). Sequence analysis was as described in section 2.4 and included confirming in-frame insertion of the sense strand sequence into the pET-21a(+) vector for protein expression in bacteria.

**Table 2.**
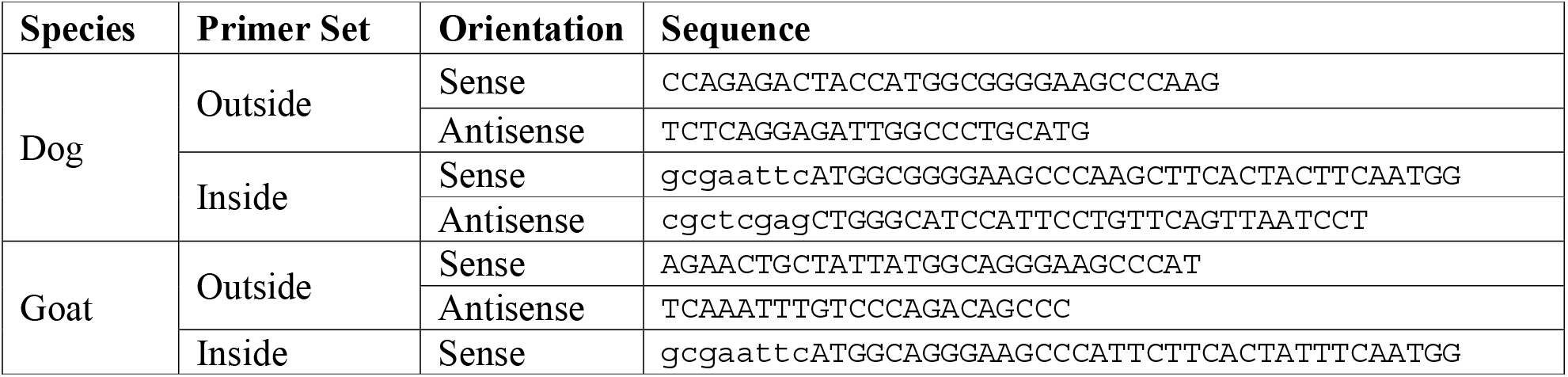
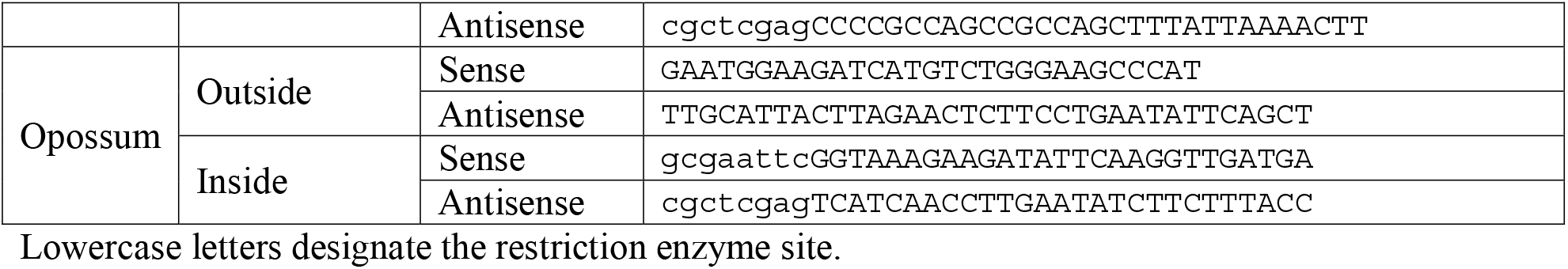
PCR primer sequences (5’–3’) for cloning GSTA3 cDNAs into pET-21a+.

## 3. Results

### 3.1. Alignment of the cloned GSTA3 mRNA sequences to those of human and horse GSTA3 mRNAs

Fig. 2 shows the alignments of the newly cloned dog, goat, and gray short-tailed opossum *GSTA3* mRNA sequences to the known *GSTA3* mRNA sequences of human (GenBank accession number **<u>NM_000847.4</u>**) and horse (**<u>KC512384.1</u>**). The 5’ ends of the sequences begin with the start codons (indicated by green font), and the 3’ ends terminate with stop codons (in red font). As expected, there is a high degree of sequence conservation across the entire mRNA sequences (gray highlighted sequences in Fig. 2 are identical to those of human *GSTA3* mRNA). The GenBank accession numbers of the newly cloned *GSTA3* cDNAs are **<u>KJ766127</u>** (dog), **<u>KM578828</u>**(goat), and **<u>KP686394</u>**(gray short-tailed opossum). The overall percentages of identical residues at each position were compared pairwise between each *GSTA3* mRNA, and are presented in Table 3. Compared to the human *GSTA3* mRNA, the dog *GSTA3* mRNA has the highest sequence identity to that of the human, with 88% identical bases. Goat *GSTA3* mRNA was similar to that of the horse in having 84% and 85% identical residues, respectively, compared to the human *GSTA3* mRNA. The most divergent was the gray short-tailed opossum *GSTA3* mRNA, which had identities to the four eutherian *GSTA3* mRNAs ranging from 68% to 71% identical bases.

**Table 3.** The percentages of identical nucleotides at each position between GSTA3 mRNAs of different mammalian species.

|  | **Human** | **Dog** | **Horse** | **Goat** | **Opossum** |
| --- | --- | --- | --- | --- | --- |
| **Human** | 100 | 88 | 85 | 84 | 70 |
| **Dog** | 88 | 100 | 87 | 87 | 71 |
| **Horse** | 85 | 87 | 100 | 85 | 70 |
| **Goat** | 84 | 87 | 85 | 100 | 68 |
| **Opossum** | 70 | 71 | 70 | 68 | 100 |

**Figure 2.**
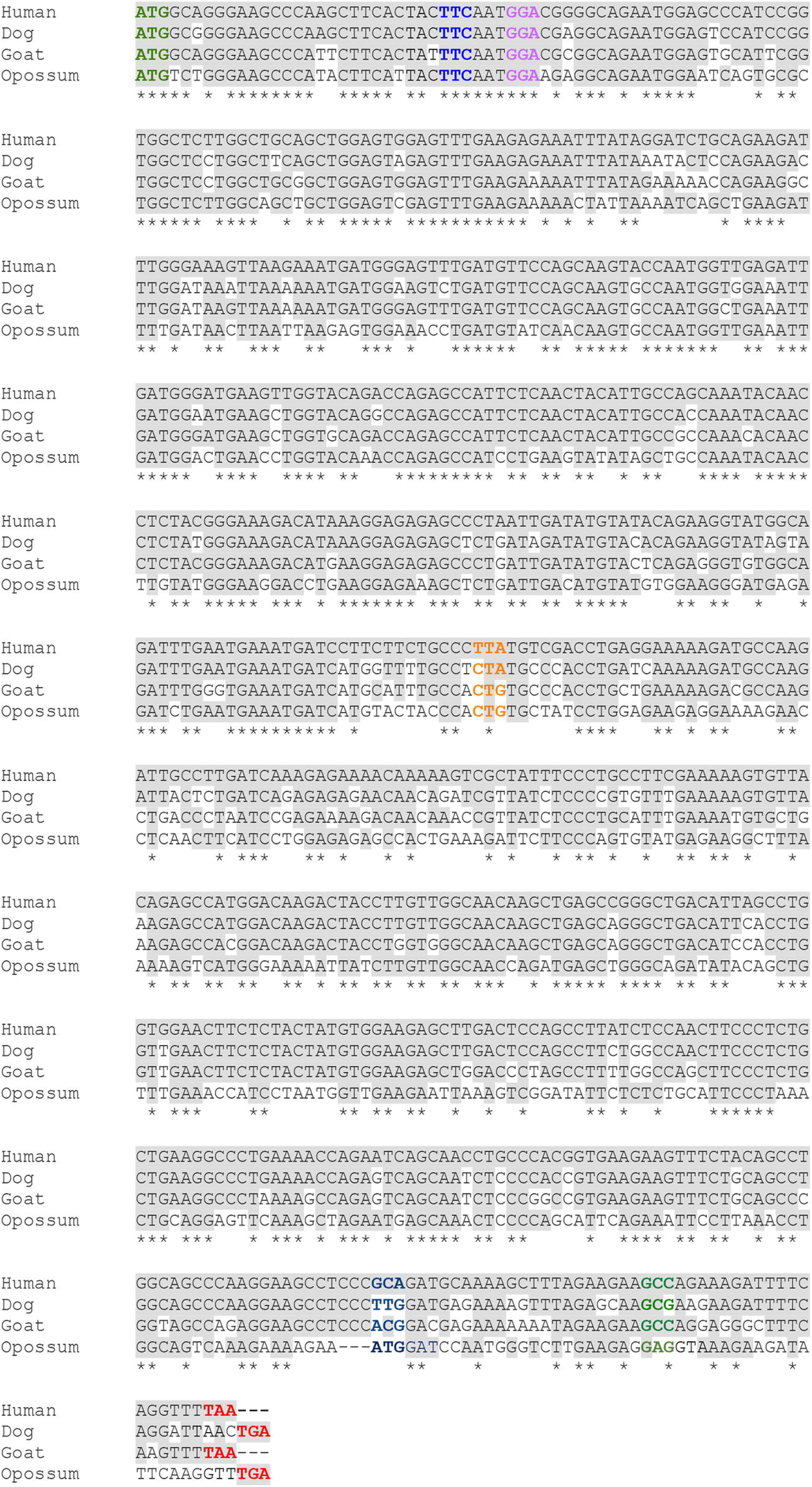
Aligned nucleotide sequences of the coding sequences of GSTA3 mRNAs from selected species including those used as reference. Human (*Homo sapiens;* **<u>NM_000847.5</u>**), horse (*Equus caballus;* **<u>KC512384.1</u>**), dog (*Canis lupus familiaris;* **<u>KJ766127</u>**), goat (*Capra hircus;* **<u>KM578828</u>**), and opossum (*Monodelphis domestica;* **<u>KP686394</u>**) those displayed here. Dog, goat, and opossum cDNA sequences were cloned in the current study. Shared identities with the human reference sequence are highlighted in gray. Start and stop codons are identified in bold, green and red type, respectively. The five critical codons defining high ketosteroid isomerase activity in human GSTA3-3 as compared to the low activity of human GST A2-2 [9] are identified in bold, multicolored type. Each full row has 60 nucleotides.

### 3.2. Alignment of the predicted GSTA3 protein sequences from five mammals

The amino acid sequences of the GSTA3 proteins were inferred from the cloned *GSTA3* mRNA sequences. They are aligned with the human and horse GSTA3 protein sequences (GenBank accession numbers **<u>NP_000838.3</u>** and **<u>AGK36275.1</u>**, respectively) in Fig. 3. Underneath the alignment, the “*” symbols identify sites with identical residues across all five species; these comprise 115 residues of the 222 total amino acids (52%) of the GSTA3 proteins. The “:”symbols identify sites with highly similar residues (53 residues of 222 total amino acids, or 24%) and the “.” symbols indicate positions with less similar residues (11 residues of the whole GSTA3 protein, or 5%). The conservation of amino acid residues between GSTA3 proteins of different species appears to be fairly consistent across the length of the proteins. The percentage of identical amino acids in the various positions was calculated between pairs of aligned GSTA3 protein sequences of all species (Table 4). The degree of conservation to human GSTA3 protein was maintained within eutherian species, with the dog GSTA3 protein having 85% identical residues, and horse and goat GSTA3 proteins both had 81% identical residues compared to the human protein. The metatherian gray short-tailed opossum GSTA3 protein had the lowest conservation with 64% identical residues with the human protein.

**Table 4.** The percentages of identical amino acids at each position between GSTA3 proteins of different mammalian species.

|  | Human | Dog | Horse | Goat | Opossum |
| --- | --- | --- | --- | --- | --- |
| Human | 100 | 85 | 81 | 81 | 64 |
| Dog | 85 | 100 | 81 | 83 | 62 |
| Horse | 81 | 81 | 100 | 82 | 61 |
| Goat | 81 | 83 | 82 | 100 | 62 |
| Opossum | 64 | 62 | 61 | 62 | 100 |

**Figure 3.**
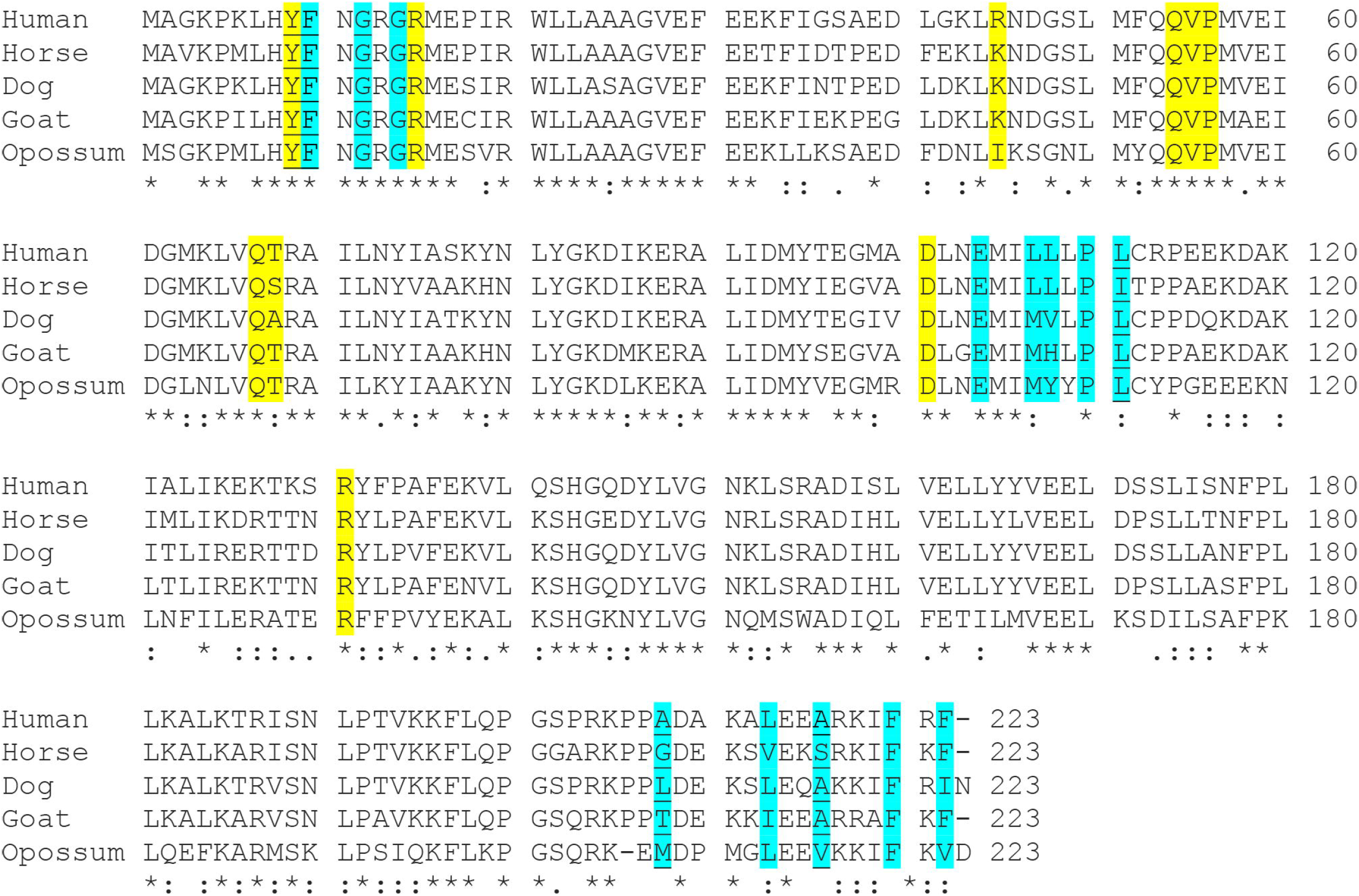
Aligned amino acid sequences for GSTA3 of all species investigated including those used as reference. Symbols: * = identical residues, : = similar residues, . = conservation of groups with weakly similar properties. G-site residues are highlighted in yellow and H-site residues are highlighted in blue. Phenylalanine in position 220 may contribute to both G- and H-sites. The underlined amino acids govern high or low 3-ketosteroid isomerase activity between the human GST A3-3 and GST A2-2.

### 3.3. Conservation of amino acids critical for steroid isomerase activity and binding of the cofactor glutathione

The G-site residues for binding the cofactor glutathione are Tyr9, Arg15, Arg45, Gln54, Val55, Pro56, Gln67, Thr68, Asp101, Arg131 and Phe220 in human GSTA3 [15]. The G-site residues were identical or conservatively changed among the GSTA3 proteins of the five species with the exception of Arg45 or Lys45 being Ile45 in the opossum enzyme (Fig. 3). Most of the H-site residues are nearly identical across all of the species. The five critical amino acids defining high ketosteroid isomerase activity in human GSTA3-3 as compared to the low activity of human GST A2-2 [9] are marked with underlining in Fig. 3 and “*” in Table 5, and are identical or similar to these human GST A3-3 residues in the three new sequences, with the exception of more bulky residues in position 208. Table 5 shows the H-site residues in the GSTA enzymes expressed in pig (“GSTA2-2”) and bovine (“GSTA1-1”) steroidogenic tissues and compared for ketosteroid isomerase activities to equine and human GSTA3-3 enzymes [15]. Both pig GSTA2-2 and bovine GSTA1-1 enzymes are similar to the dog, goat and opossum enzymes in having have more bulky residues at position 208 than do the human and equine GSTA3-3 enzymes. This may be part of the reason that the pig and bovine enzymes are only 0.01-to 0.003-fold as active with the substrate Δ^5^-androstene-3,17-dione compared to the equine GSTA3-3 enzyme. The divergent residues in positions 107 and 108 of the dog, goat, opossum GSTA3 and the bovine GSTA1 proteins are probably also relevant to isomerase activities.

**Table 5.**
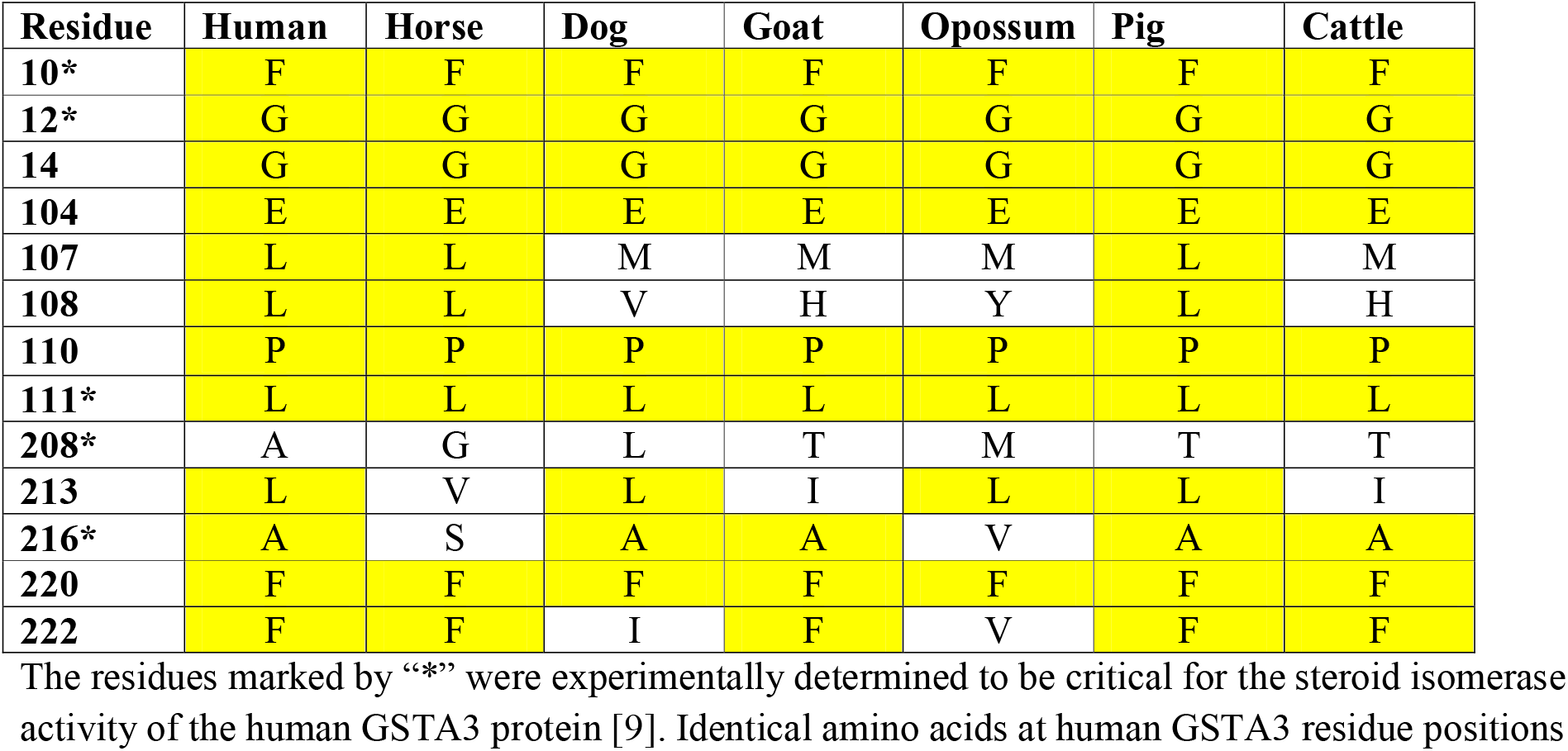
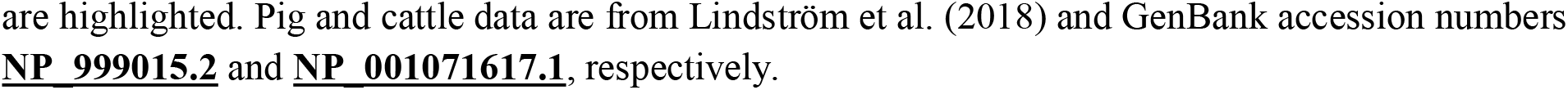
Conservation of H-site residues involved in steroid isomerization across GSTA3 proteins of different mammalian species.

| Residue | Human | Horse | Dog | Goat | Opossum | Pig | Cattle |
| --- | --- | --- | --- | --- | --- | --- | --- |
| **10\*** | F | F | F | F | F | F | F |
| **12\*** | G | G | G | G | G | G | G |
| **14** | G | G | G | G | G | G | G |
| **104** | E | E | E | E | E | E | E |
| **107** | L | L | M | M | M | L | M |
| **108** | L | L | V | H | Y | L | H |
| **110** | P | P | P | P | P | P | P |
| **111\*** | L | L | L | L | L | L | L |
| **208\*** | A | G | L | T | M | T | T |
| **213** | L | V | L | I | L | L | I |
| **216\*** | A | S | A | A | V | A | A |
| **220** | F | F | F | F | F | F | F |
| **222** | F | F | I | F | V | F | F |

The residues marked by “\*” were experimentally determined to be critical for the steroid isomerase activity of the human GSTA3 protein [9]. Identical amino acids at human GSTA3 residue positions
are highlighted. Pig and cattle data are from Lindström et al. (2018) and GenBank accession numbers **NP\_999015.2** and **NP\_001071617.1**, respectively.

### 3.4. Modeling the structures of the new GST A3-3

The crystal structure of human GST A3-3 in complex with Δ^4^-androstene-3,17-dione has been determined [22] and was used to model the structures of the related novel GST proteins. All GST A3-3 proteins appear to fold in a similar manner, and possibly could accommodate the steroid substrate in a productive binding mode without major clashes with the H-site residues. The dog GST model shows favorable interactions with the steroid except the close approach of Leu 208 to the D-ring of the substrate (Fig. 4). The high-activity human and equine GST A3-3 enzymes have smaller residues in position 208, Ala208 and Gly208, respectively. Similarly, the modeled opossum GST A3-3 enzyme indicates that its Met208 overlaps with the steroid, unless the overlap can be avoided by a rotation of the Met sidechain. The goat GST A3-3 has Thr208 in this highly variable position, and this residue is smaller than Leu208 in the dog enzyme. On the basis of the modeling, the dog, opossum, and goat GST A3-3 enzymes might have steroid isomerase activity, based on comparison with the structure of the high-activity human GST A3-3 in complex with androstenedione [22], but their more bulky 208 residues is a caveat.

**Figure 4.**
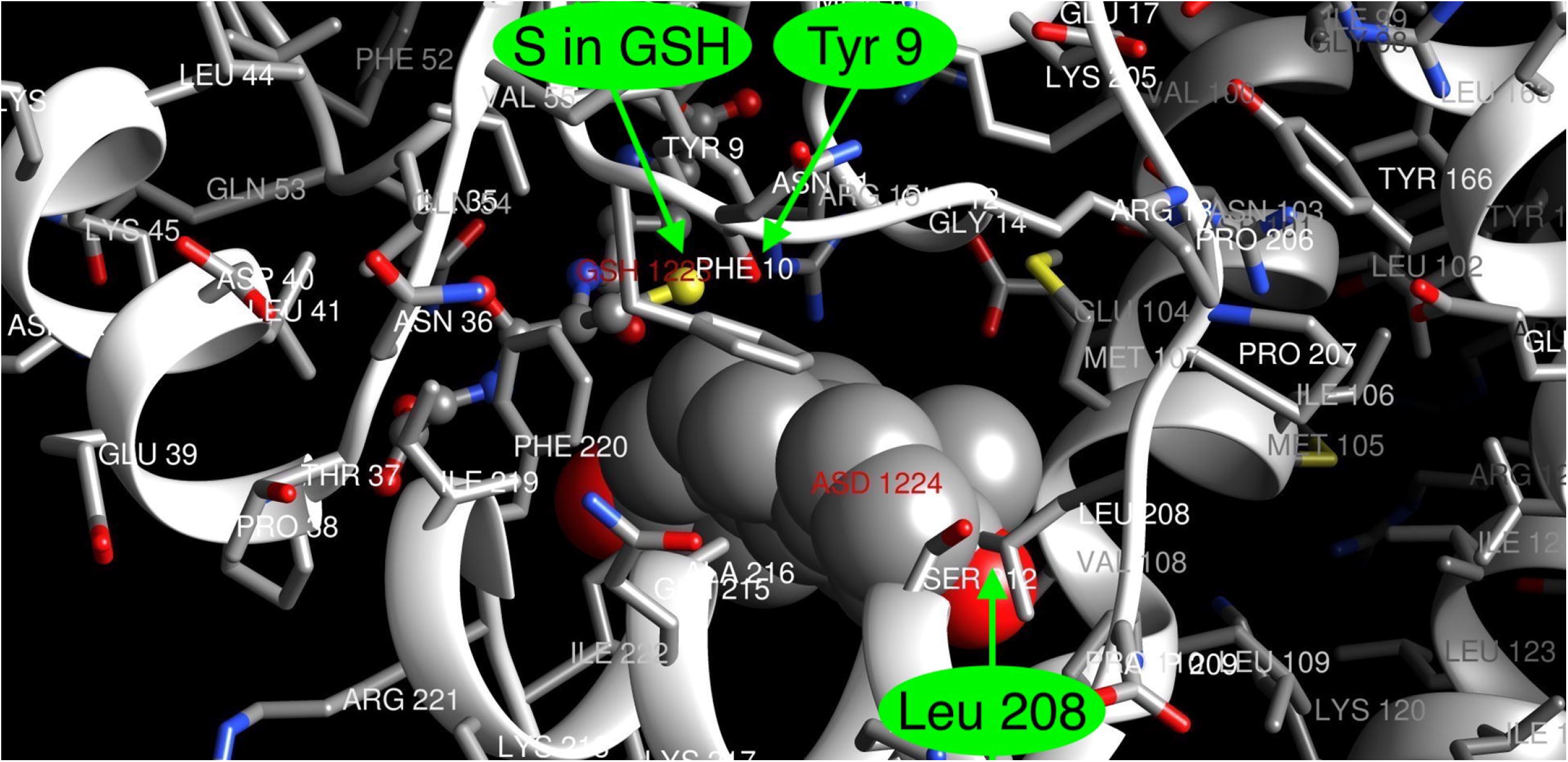
Dog GST A3-3 active site modeled with bound Δ^4^-androsten-3,17-dione. The crystal structure of the human GST A3-3 with the ligands glutathione and the steroid (PDB code 2vcv) was used as a template for modeling the dog enzyme. The ligands in the dog GST A3-3 were positioned by superpositioning the human GST A3-3 onto the dog model. Green arrows indicate the location of the residue Leu208 very close to the D-ring of Δ^4^-androsten-3,17-dione, as well as the sulfur of glutathione and the oxygen of Tyr9 implicated in the catalysis of steroid isomerization. The image was created with the UCSF Chimera package (http://www.rbvi.ucsf.edu/chimera) developed by the Resource for Biocomputing, Visualization, and Informatics at the University of California, San Francisco (supported by NIGMS P41-GM103311).

## 4. Discussion

Complementary DNAs were cloned from *GSTA3* mRNAs in testes from the dog, goat, and gray short-tailed opossum. Given that the sequences of the *GSTA1, GSTA2* and *GSTA3* mRNAs are highly conserved [10], cloning of *GSTA3* mRNA was difficult even from testes, which has a distinctively high level of *GSTA3* gene expression [15]. The PCR primers used here were designed to have their 3’ ends bind to codons or untranslated sequences that are unique to the *GSTA3* mRNA. Interestingly, there are no reagents for nucleic acid hybridization or antibody immunodetection that can distinguish *GSTA3* gene products from those of *GSTA1* and *GSTA2* genes, both of which are ubiquitously expressed. The *GSTA3* mRNA can only be specifically amplified using PCR primers, such as those designed here and previously [11,18]. For example, in order to localize *GSTA3* gene expression to Leydig cells within horse testes, we developed an *in situ* reverse transcription-PCR protocol that was specific for *GSTA3* mRNA [18].

The human GSTA3-3 enzyme has well characterized catalytic efficiency with substrates Δ^5^–androstene-3,17-dione and Δ^5^ – pregnene-3,20-dione, which is second only to the equine GSTA3 enzyme of the six GSTA proteins characterized so far [15]. The H-site residues of the GSTA3 proteins are critical for positioning the steroid substrate to be isomerized. The amino acids required for that activity were initially determined experimentally for human GSTA3-3 [9,21]. The human and horse GSTA3 proteins have very high conservation of H-site residues (Table 5). These differ from related GSTA enzymes, such as human GSTA2-2, which have very little activity [15]. When activity data become available for the dog, goat, and opossum enzymes, we may be able to relate enzymatic activity differences to specific amino acids or combinations of them. It will also be instructive to compare the steroid isomerase activities of the newly cloned GSTA3-3 enzymes with those of the bovine [23] and pig [10] cognates of the human GSTA3-3, both of which have significantly lower steroid isomerase activities than the human and horse GSTA3-3 enzymes [15].

As a recently discovered enzyme in steroid hormone biosynthesis, the GSTA3-3 enzyme and its corresponding mRNA sequence require extensive functional analyses across species. The mechanism of GSTA3-3 action and regulation of the *GSTA3* gene must be identified in order to fully understand the role played in steroid hormone biosynthesis and the possible implications of the enzyme function being impaired. Our work can contribute through identification of the *GSTA3* mRNA coding sequence in multiple species, as well as providing clones in the expression vector pET-21a(+) which can produce GST A3-3 proteins to be evaluated for their activities as steroid isomerases. Evaluation of these recombinant enzymes will enable quantitative comparisons of the steroid isomerase activities among the species investigated, and allow further investigation of the effects of endogenous compounds like follicle stimulating hormone, luteinizing hormone, testosterone, estradiol, and glucocorticoids, as well as pharmaceuticals like phenobarbital or dexamethasone on the expression of the enzyme. For instance, expression of the *GSTA3* gene was down-regulated along with testosterone synthesis in testes of stallions treated with dexamethasone [18,19]. In cultured porcine Sertoli cells, follicle stimulating hormone, testosterone and estradiol increased *GSTA* gene expression [24]. In addition, inhibitors of the GSTA3-3 enzyme have been identified that may be medically useful for reducing fertility or progression of steroid-dependent diseases [15].

## 5. Conclusion

The three new *GSTA3* mRNA and protein sequences we report extend our comparative knowledge of the GST A3-3 enzymes of diverse mammalian species; and the information and reagents we have generated will help to facilitate future studies to characterize the GST A3-3 enzymes more generally. In addition, they may augment studies of the regulation of the expression of *GSTA3* genes in steroidogenic and other tissues. Mechanistic data of this kind can enable development of new approaches for manipulating GSTA3-3 enzyme activities *in vivo* for medical purposes. These could include inducing interruptions in fertility in animal species important to man or developing novel pharmacological treatments for steroid-dependent diseases in man and animals. It is essential that the new treatments that are prescribed are both effective and specific. It is, therefore, crucial that researchers continue to investigate steroidogenesis and its regulation in order to generate more complete mechanistic knowledge.

## Acknowledgments

The authors acknowledge the assistance of Dr. John Edwards with tissue collection. Additionally, B.M. was supported by the Swedish Research Council (grant 2015-04222).

